# Increase in weighting of vision vs. proprioception associated with force field adaptation

**DOI:** 10.1101/544189

**Authors:** Brandon M. Sexton, Yang Liu, Hannah J. Block

## Abstract

Hand position can be encoded by vision, via an image on the retina, and proprioception (position sense), via sensors in the joints and muscles. The brain is thought to weight and combine available sensory estimates to form an integrated multisensory estimate of hand position with which to guide movement. Force field adaptation, a form of cerebellum-dependent motor learning in which reaches are systematically adjusted to compensate for a somatosensory perturbation, is associated with both motor and proprioceptive changes. The cerebellum has connections with parietal regions thought to be involved in multisensory integration; however, it is unknown if force adaptation is associated with changes in multisensory perception. One possibility is that force adaptation affects all relevant sensory modalities similarly, such that the brain’s weighting of vision vs. proprioception is maintained. Alternatively, the somatosensory perturbation might be interpreted as proprioceptive unreliability, resulting in vision being up-weighted relative to proprioception. We assessed visuo-proprioceptive weighting with a perceptual estimation task before and after subjects performed straight-ahead reaches grasping a robotic manipulandum. Each subject performed one session with a clockwise or counter-clockwise velocity-dependent force field, and one session in a null field to control for perceptual changes not specific to force adaptation. Subjects increased their weight of vision vs. proprioception in the force field session relative to the null field session, regardless of force field direction, in the straight-ahead dimension (F_1,44_ = 5.13, p = 0.029). This suggests that force field adaptation is associated with an increase in the brain’s weighting of vision vs. proprioception.

## Introduction

To keep voluntary movement accurate in the face of internal or environmental perturbations, the brain may make adjustments in both sensory and motor systems. In the context of motor learning, sensory changes have been suggested in both animal (Xerri et al., 1999) and human studies (David J. Ostry et al., 2010). Xerri et al. (1999) trained monkeys to pick up food pellets. As the monkeys learned the task, they began using smaller regions of their fingers to pick up the pellets. The corresponding representations in somatosensory cortex (S1) grew about two times larger than the same S1 fingertip regions for the contralateral hand, suggesting an effect of motor learning on the somatosensory system (Xerri et al., 1999). Ostry et al. (2010) instructed human subjects to make straight-ahead reaches while grasping a robotic manipulandum that applied a velocity-dependent force field. Initial movement errors were reduced with trial-and-error practice, a sign of cerebellum-dependent motor adaptation (Baizer et al., 1999; Block and Bastian, 2012; T.A. Martin et al., 1996); interestingly, Ostry et al. (2010) found systematic changes in somatosensation (kinesthesia) in the adapted arm. These perceptual changes may indicate the involvement of somatosensory cortex in motor adaptation (Darainy et al., 2013; Vahdat et al., 2014). In addition, visuomotor adaptation to a cursor rotation results in systematic proprioceptive changes for the adapted hand (Clayton et al., 2014; Henriques and Cressman, 2012), which is consistent with somatosensory involvement in motor learning whether the perturbation is visual (cursor rotation) or somatosensory (force field).

While somatosensory involvement in motor adaptation has been an important area of investigation in the last ten years, the potential role of multisensory processing in motor adaptation has yet to be considered. Multiple sensory systems play an important role in voluntary movement. For example, to plan an accurate reach, the brain must have an accurate initial estimate of the hand’s position. We normally have access to true hand position (*Y*) through vision and proprioception. The image of the hand on the retina provides a visual estimate (*ŷ* _*V*_), while receptors in the muscles and joints of the arm provide a proprioceptive estimate (*ŷ*_*P*_). To form a single estimate with which to guide behavior, the brain is thought to weight and combine them into a single, integrated estimate of hand position (Ghahramani et al., 1997). If *ŷ* _*VP*_ is this integrated estimate, and *W*_*V*_ is the weight of vision vs. proprioception (*W*_*V*_ < 0.5 implies greater reliance on proprioception):

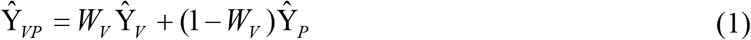

Although the neural basis of this process is unknown, it may involve multisensory regions of posterior parietal cortex such as the intraparietal sulcus (Limanowski and Blankenburg, 2016) or angular gyrus (Block et al., 2013). The weighting of each sensory input is thought to be inversely proportional to the associated variance; this is known as the minimum variance or maximum likelihood estimation model (Bays and Wolpert, 2007), and it has experimental support from a variety of human behaviors (Ernst and Banks, 2002; Ghahramani et al., 1997; Liu et al., 2018; van Beers et al., 1996).

Importantly, the reliability of a given sensory input, and its weighting in multisensory integration, is not constant, but may vary with environmental conditions (Mon-Williams et al., 1997) and locus of attention (Block and Bastian, 2010; Warren and Schmitt, 1978; Welch and Warren, 1980). Thus, multisensory integration can respond to changes in the body or environment that affect sensory perception. For example, a decrease in illumination likely results in a multisensory estimate that relies more on proprioception than vision (Mon-Williams et al., 1997). In addition, the computation being performed may affect sensory weighting; subjects rely more on vision for planning movement vectors and more on proprioception for planning the joint-based motor command (Sober and Sabes, 2003). The modality of the target being reached also plays a role, with subjects minimizing coordinate transformations by relying more on vision when reaching to visual targets, and more on proprioception when reaching to proprioceptive targets (Sober and Sabes, 2005). Even different aspects of what might be considered the same computation can use different weightings; vision is weighted more heavily relative to proprioception when localizing the hand in azimuth than in depth (van Beers et al., 2002).

Most of the evidence of multisensory involvement in motor learning comes from studies asking which sensory signals are necessary for force field adaptation. This form of motor adaptation can occur without a proprioceptive error, using visual feedback (Melendez-Calderon et al., 2011; Miall et al., 2018; Sarlegna et al., 2010), but also without a visual error, using only proprioception (Miall et al., 2018; Scheidt et al., 2005). This suggests subjects can flexibly use available error information, whether visual or proprioceptive (Miall et al., 2018). Results from Haith et al. (2008) suggest spatial recalibration of both visual and proprioceptive estimates occurs after force field learning (Haith et al., 2008). However, simultaneous visual and proprioceptive processing, where visuo-proprioceptive integration can occur, has not been considered in the context of force adaptation.

Here we asked whether force field adaptation affects the brain’s weighting of visual and proprioceptive estimates of hand position when both are available. Given that multisensory integration plays a key role in movement planning, one possibility is that force field adaptation affects all relevant sensory modalities similarly, such that a constant weighting of vision vs. proprioception is maintained. Alternatively, the somatosensory perturbation could be considered a source of proprioceptive unreliability, which we would expect to result in vision being up-weighted relative to proprioception. We assessed visuo-proprioceptive weighting with a perceptual estimation task before and after subjects performed straight-ahead reaches while grasping a robotic manipulandum. Each subject performed one session with reaches in a clockwise or counter-clockwise velocity-dependent force field, and one session in a null field to control for perceptual changes not specific to force adaptation.

## Methods

### Subjects

46 healthy right-handed adults (aged 18-33, mean age 22.7 years; 22 female) completed two sessions each, scheduled at least 4 days apart. Subjects reported that they were free of neurological or musculoskeletal problems, and had normal or corrected-to-normal vision in both eyes. All procedures were approved by the Indiana University Institutional Review Board, and subjects provided written informed consent.

### Session design

Subjects were randomly assigned to the clockwise (CW) or counterclockwise (CCW) group (N=23 each), and to have the real or null session first. In each session, subjects performed a series of straight-ahead reaching movements grasping a robotic manipulandum (KINARM End Point, BKIN), with a sensory estimation task to assess visual and proprioceptive perception of hand position before and after the reaching task. In the real session, subjects adapted to a CW or CCW velocity-dependent force field, according to group assignment. The null session comprised the same number of reaching movements, but no force field. This was intended to control for any perceptual changes not specific to force field adaptation, such as making reaching movements, grasping the manipulandum, viewing the task display, etc.

Each session consisted of five blocks of trials (Fig. 1A): baseline reaching in the null field with the left and right hands (16 trials each hand); pre-adaptation visuo-proprioceptive estimation task (35 trials); adaptation block of reaching with the right hand (208 trials with the right hand in a real or null field depending on session); post-adaptation visuo-proprioceptive estimation task (35 trials); and washout reaching in the null field with the left and right hands (16 trials each hand). Left hand baseline and washout reaching trials were included so we could assess any intermanual transfer of force field learning, as the left hand was used to indicate the subject’s perception of right hand position in the visuo-proprioceptive estimation task.

**Figure 1.**
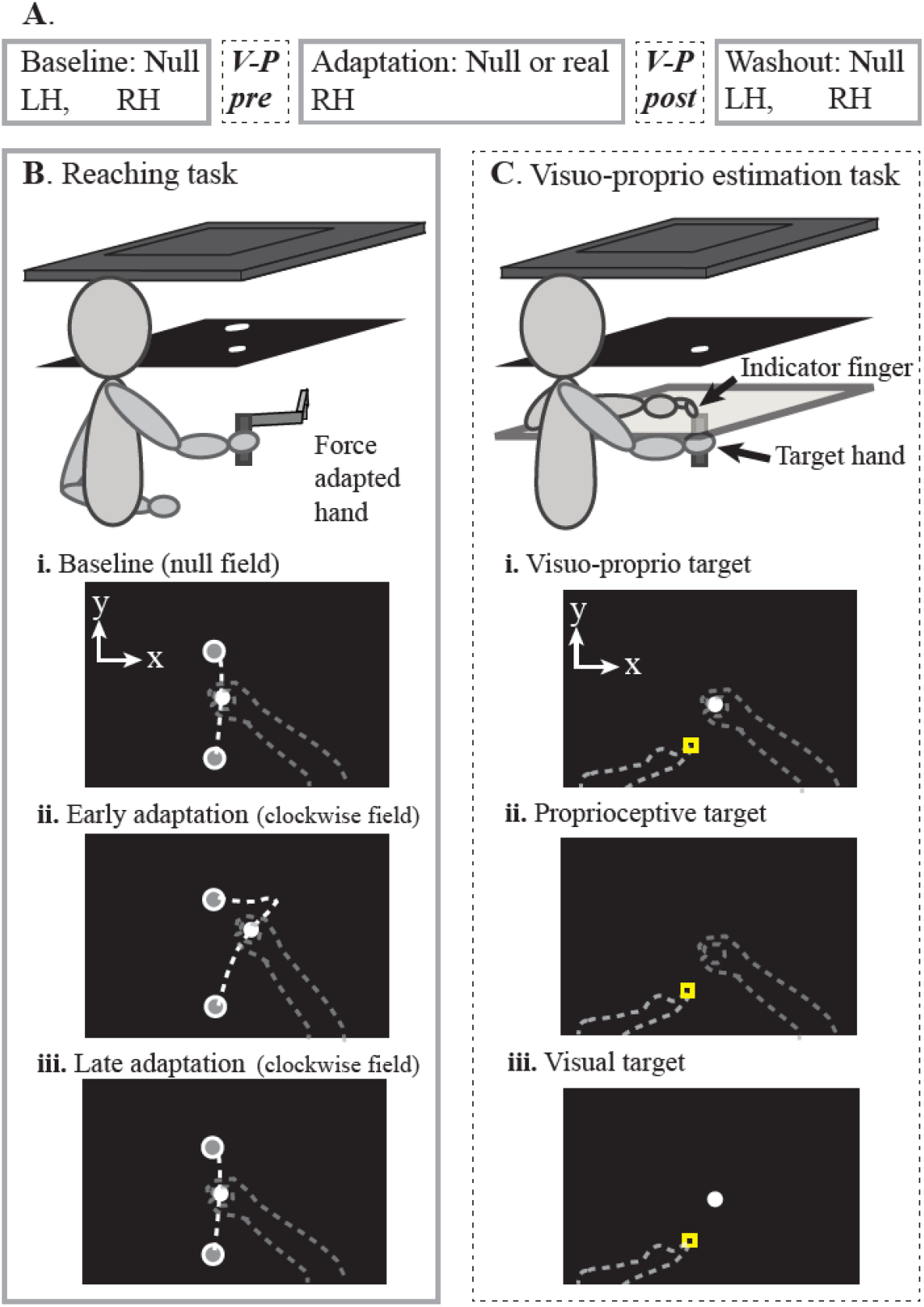
**A.** Single session protocol. After practicing each task, subjects completed five blocks of reaching (solid outline) and two blocks of visual and proprioceptive estimates (dashed outline). Each subject performed two sessions, one of which included a velocity-dependent force field during the adaptation block. Subjects had no direct vision of either hand at any point in the session. **B. Reaching task**. Subjects viewed images displayed on a TV (top) in a horizontal mirror. Subjects were instructed to make a series of straight-ahead movements grasping a robotic manipulandum. Images in the mirror appeared to be in the plane of the manipulandum, which was represented continuously by a white cursor. **i.** Top-down view of the display. Dashed lines not visible to subject. During baseline and washout blocks of both sessions, and the adaptation block of the null session, subjects moved the passive manipulandum (null field) from the start circle to the target circle, both at body midline. **ii.** A CW or CCW field was applied during the adaptation block of the real session only. Reaching paths early in this block had large rightward or leftward perpendicular deviations, respectively. **iii.** After subjects had adapted to the force field, reaching trajectories became straight, with perpendicular deviations similar to baseline. **C. Visuo-proprioceptive estimation task**. At the same apparatus, a touchscreen was placed directly over the manipulandum. Subjects moved their left (indicator) index finger, on top of the touchscreen, from a variable start position (square) to the perceived location of targets related to the right hand (target hand), which was below the touchscreen. Subjects were instructed to take their time and place their indicator finger as accurately as possible. No online or endpoint feedback about the indicator finger was given. **i.** For the VP target, a white disc appeared above the center of the manipulandum handle, grasped in the right hand. **ii.** The P target was the same except no white disc was displayed. **iii.** The V target was the white disc alone, with the right hand resting in the subject’s lap.

### Apparatus

Subjects were seated in front of a reflected rear projection apparatus (TV and mirror) throughout the session, such that the task display appeared to be in the same horizontal plane as the manipulandum (Fig. 1). Both hands remained below the mirror at all times, preventing direct vision of either hand, and a cape attached to the edge of the mirror was draped over subjects’ shoulders to prevent vision of the upper arms. Subjects were centered with the apparatus and strapped to the chair with a harness to limit torso motion. A headband across the forehead was attached to the edge of the TV with Velcro, restricting subjects’ head motion.

### Reaching task

Subjects grasped the manipulandum handle and made a series of straight-ahead movements to a visual target 20 cm from the start position (Fig. 1B). A 1cm white disc was displayed over the center of the manipulandum handle, providing online feedback throughout the reach. Subjects were instructed to make their movement paths straight and brisk. During the adaptation block of the real force session only, subjects experienced a CW or CCW velocity-dependent force field (Fig. 1Bii):

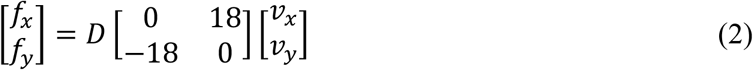

where *f*_*x*_ and *f*_*y*_ are the commanded force to the manipulandum in the lateral (x) and sagittal (y) directions, *v*_*x*_ and *v*_*y*_ are hand velocities, and *D* is the force direction (1 for CW, −1 for CCW). The desired movement time was 575-650 ms, and subjects received feedback telling them when movements were too fast or too slow. Each trial ended when the manipulandum reached the target, at which point the hand was passively moved back to the start position. The maximum perpendicular deviation of the manipulandum from a straight-line path was computed for each trial as a measure of movement error. To normalize movement errors to baseline levels, max perpendicular deviation at the end of right hand baseline was subtracted from every right hand trial, and max perpendicular deviation at the end of left hand baseline was subtracted from every left hand trial.

To quantify the degree of perturbation at the beginning of the adaptation block, we averaged max perpendicular error on the first 8 trials of adaptation. The washout blocks were used to estimate negative aftereffect (1) in the right hand as a measure of force field learning, and (2) in the left hand to measure transfer of learning to the untrained left hand, which was used to indicate the subject’s perceptions in the sensory estimation task. Negative aftereffect was estimated by taking the mean of the first 8 washout trials in each hand.

### Visuo-proprioceptive estimation task

Immediately before and after the force adaptation block, subjects performed a sensory estimation task to assess visual and proprioceptive estimates of their right hand position (Fig. 1C). With no direct vision of either hand, subjects used their left (indicator) index finger to point to the perceived location of three target types (Block and Bastian, 2011, 2010; Liu et al., 2018; Munoz-Rubke et al., 2017): a visuo-proprioceptive target (1 cm white disc displayed directly above their right hand, which grasped a stationary manipulandum handle beneath the touchscreen glass) (Fig. 1 Ci); proprioceptive-only target (right hand grasping the handle, with no white disc) (Fig. 1 Cii); and a visual-only target (white disc alone, with the right hand lowered to rest in the subject’s lap) (Fig. 1 Ciii). For this task, a touchscreen, consisting of a 3-mm pane of glass with an infrared touch overlay (PQ Labs), was slid into place directly above the robotic manipulandum (and below the mirror) to record indicator finger positions. To prevent subjects from learning or repeating a particular movement direction or extent with the left hand, indicator finger start position was randomized trial-to-trial between five start positions, and targets between two target positions, all centered with the body midline. Subjects received step-by-step instructions to guide them through the task via pre-recorded audio prompts. Subjects received no online or endpoint feedback about the left indicator finger and no knowledge of results. There were no speed requirements, and subjects were instructed to take their time and be as accurate as possible. Subjects were asked not to slide their finger on the glass. Adjustments of indicator finger position were permitted, with the final position recorded once the finger had not moved more than 2 mm in 2 seconds.

Each block of the visuo-proprioceptive estimation task (pre-and post-adaptation) comprised 35 trials: 15 visual-only (V), 15 proprioceptive-only (P), and 5 visuo-proprioceptive (VP) trials, in pseudorandom order. We computed an estimate of subjects’ weighting of vision vs. proprioception (*wv*) when both modalities were available, on VP targets (Block and Bastian, 2010). Because *wv* has been observed to differ across spatial dimensions (Bays and Wolpert, 2007; van Beers et al., 2002, 1996), we computed separate values for the x-and y-dimensions by dividing the distance between P and VP target estimates by the sum of the P-to-VP and V-to-VP distances. For the x-dimension (Fig. 2A):

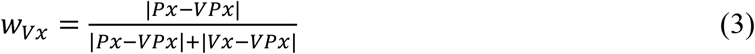

where *|Px - VPx|* and *|Vx – VPx|* are the x-dimension distances between the mean final position of the indicator finger on P or V targets, respectively, and the mean position of the indicator finger on VP targets. Similarly, for the y-dimension (Fig. 2B):

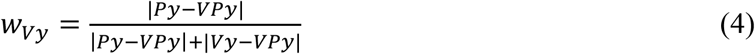

**Figure 2.**
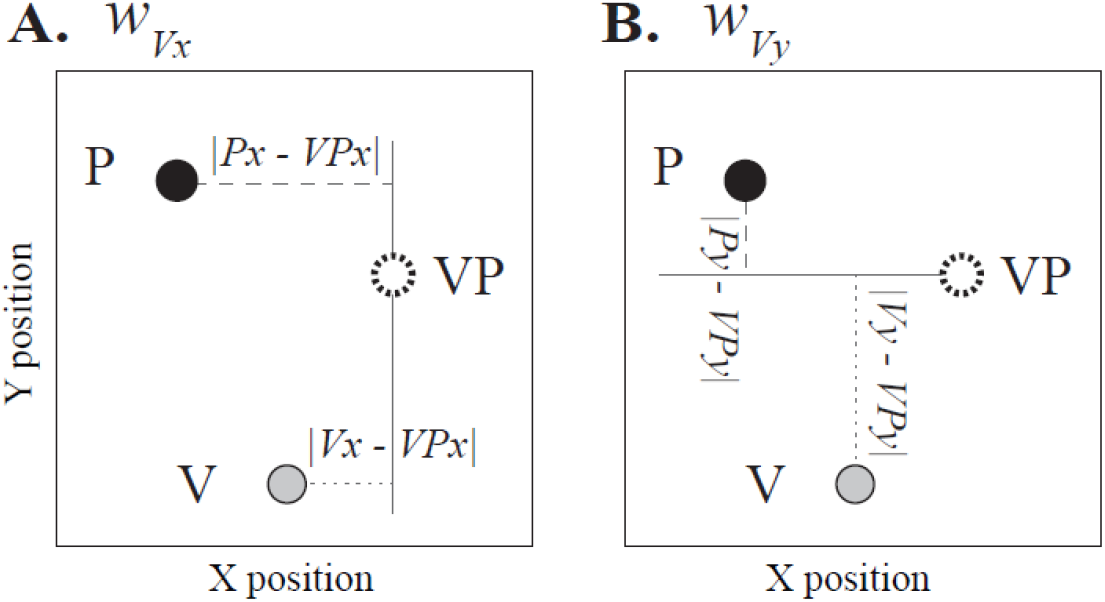
Computing weight of vision vs. proprioception. In this diagram, the subject’s estimate of V, P, and VP target positions are represented by a grey, black, and white disc, respectively. **A.** *wv*_*x*_ is computed by dividing the P-to-VP distance in the x-dimension (dashed line) by the sum of the P-to-VP distance and the V-to-VP distance in the x-dimension (dashed and dotted lines). **B.** *wv*_*y*_ is computed similarly, with all distances in the y-dimension.

In other words, if VP endpoint positions are closer to P than V positions in the y-dimension, the subject relied more on proprioception than vision (*wv*_*y*_ < 0.5). This method takes advantage of the different spatial biases inherent in vision and proprioception (Crowe et al., 1987; Foley and Held, 1972), even with no perturbation (Smeets et al., 2006). Because *wv* undergoes small fluctuations over time, as we have done previously (Block and Bastian, 2010), we computed a separate *wv* for each of the 5 VP trials pre-and post-adaptation, comparing each VP trial indicator finger endpoint with the means of the 4 V and 4 P trials occurring closest in time. The 5 values were then averaged for each subject to give a weighting estimate for the pre-or post-adaptation sensory task.

### Statistical analysis

All statistical inferences were performed two-tailed, with α of 0.05. To evaluate whether either hand experienced aftereffects after force field adaptation, we performed a separate 7paired-sample t-test for each hand in each group. In each case, we compared max perpendicular error in the appropriate washout block (right or left hand) across the real and null sessions. A significant difference between sessions would suggest the presence of aftereffect for that hand. We predicted aftereffects in the right hand, which was exposed to the force field, but not in the left hand, which was not exposed.

Separate mixed model ANOVAs (timepoint × session × group) were used to analyze *wv*_*x*_ and *wv*_*y*_. Timepoint (pre-and post-adaptation) and session (real and null) were within-subjects factors, and group (CW and CCW force field) was a between-subjects factor. A significant 3-way interaction (timepoint × session × group) would indicate that the variable changed differently in the two sessions, and that force field direction matters. In the absence of the 3-way interaction, a significant timepoint × session interaction would indicate that the variable changed differently in the two sessions, but force field direction (i.e., group) does not make a difference.

## Results

All means are given with their 95% confidence intervals (CI).

### Reaching task

In the real session, both groups had small perpendicular errors during baseline. Both CW and CCW groups initially had large perpendicular errors when the force was introduced: 20 ± 9 mm rightward and 23 ± 8 mm leftward, respectively (mean ± 95% CI). In contrast, at the same point in the null session, error was only 1 ± 5 mm leftward for the CW group, and 0.5 ± 4 mm leftward for the CCW group. After 208 force field trials in the real session’s adaptation block, error returned to approximately baseline levels, suggesting adaptation to the perturbation had occurred (Fig. 3). In the null session, max perpendicular errors were small throughout the reaching task for both groups, as expected, although blocks performed with the left hand appeared to have slight rightward errors compared to blocks performed with the right hand (Fig. 3).

**Figure 3.**
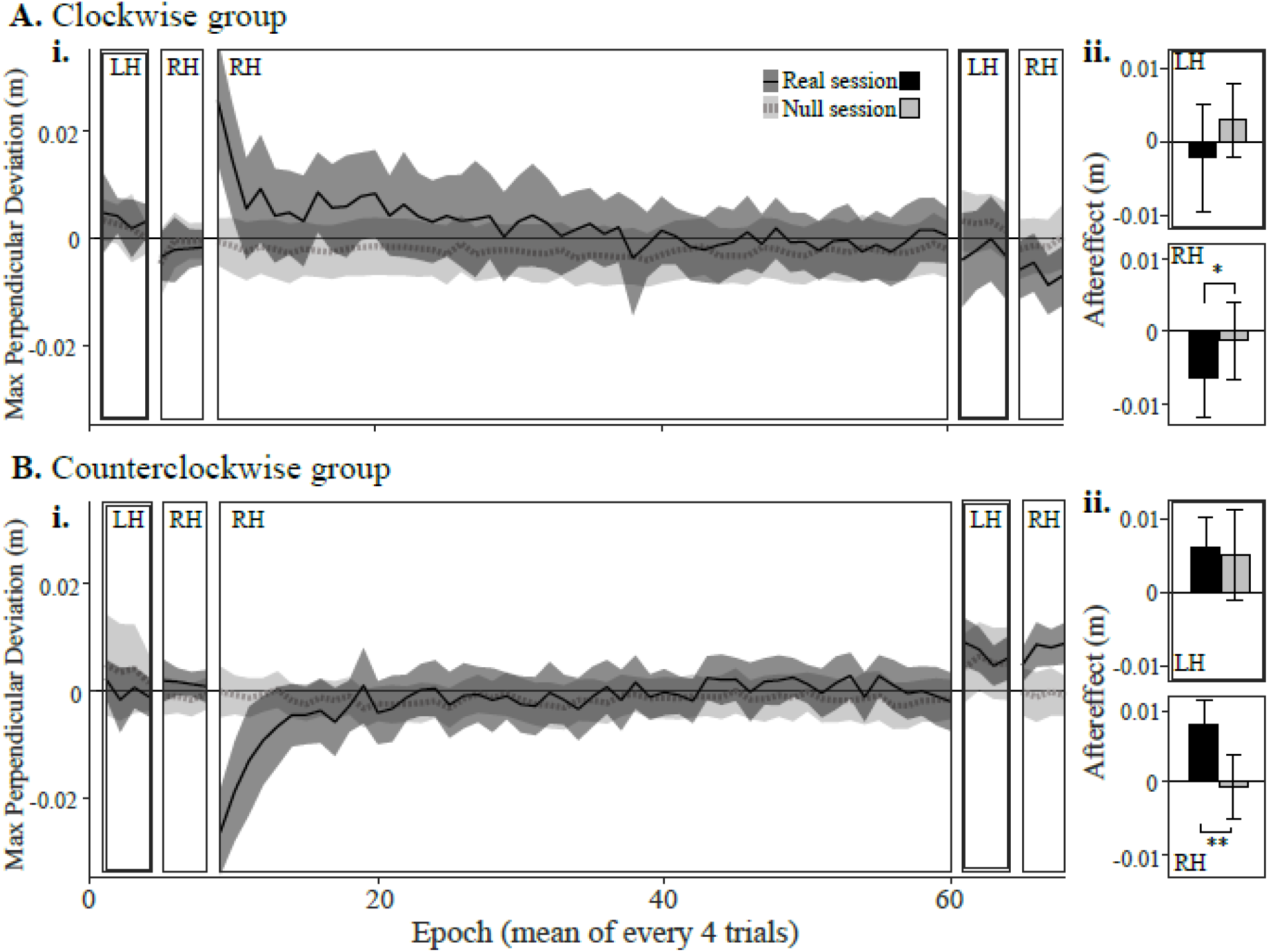
Reaching errors for the clockwise (**A**) and counterclockwise (**B**) groups. Positive values reflect rightward errors. **i.** Max perpendicular reaching error in epochs of 4 trials, averaged across subjects with 95% confidence intervals (shaded regions). In the real session (black solid line) subjects experienced a CW or CCW force field with the right hand, after a short baseline with each hand in the null field. Initial large errors on exposure to the force field decreased to near-baseline levels by the end of the adaptation block. Finally, washout blocks in the null field were used to assess negative aftereffects of force field adaptation in each hand. **ii.** Mean aftereffect in each hand with 95% confidence intervals. Top: In the left hand, neither group showed a significant difference between real and null sessions, suggesting little or no transfer of learning to the left hand in the real session. Bottom: In the right hand, both groups showed some evidence of an aftereffect in the real relative to null session, suggesting that force field adaptation occurred robustly in the real session. * p < 0.1. ** p < 0.05.

Post-adaptation washout blocks performed with the right hand tended to have larger perpendicular errors in the real session than the null session—leftward for the CW group and rightward for the CCW group—suggesting the presence of negative aftereffect. This difference did not reach statistical significance for the CW group (t_22_ = −1.98, p = 0.06), but did for the CCW group (t_22_ = 2.69, p = 0.013). In the CW group, right hand aftereffect was 7 ± 5 mm to the left in the real session, and 1 ± 5 mm to the left in the null session (Fig. 3A). In the CCW group, right hand aftereffect was 8 ± 3 mm to the right in the real session, and 0.6 ± 4 mm to the left in the null session (Fig. 3B).

Post-adaptation washout blocks performed with the left hand were similar in the real and null sessions (Fig. 3), which is not consistent with intermanual transfer of learning. In the CW group, left hand aftereffect was 2 ± 7 mm to the left in the real session, and 3 ± 5 mm to the right in the null session (t22 = −1.02, p = 0.32). In the CCW group, left hand aftereffect was 6 ± 4 mm to the right in the real session, and 5 ± 6 mm to the right in the null session (t22 = 0.27, p = 0.79).

### Visuo-proprioceptive estimation task

The example subject in Fig. 4 increased their reliance on vision over proprioception in the y-dimension (positive Δ*wv*_*y*_) in the real (CCW field) session, but not the null session. In contrast, Δ*wv*_*x*_ was slightly negative in the real session, but positive in the null session (Fig. 4). At the group level, we found Δ*wv*_*y*_ to be consistently more positive in the real session than the null session, for both groups (Fig. 5). In the CW group, Δ*wv_y_* was 0.12 ± 0.10 for the real session (mean ± 95% CI) and 0.006 ± 0.094 for the null session. In the CCW group, Δ*wv*_*y*_ was 0.020 ± 0.10 for the real session and −0.075 ± 0.083 for the null session. There was no main effect of timepoint (pre, post), session (real, null) or group (CW, CCW) for *wv*_*y*_ (F_1,44_ = 0.54, 0.49, 0.32, respectively, and p = 0.46, 0.49, and 0.57, respectively). However, there was a significant interaction of timepoint × session (F_1,44_ = 5.13, p = 0.029). The interaction of timepoint × session × group was not significant (F_1,44_ = 0.08, p = 0.78), nor was the interaction of session × group (F_1,44_ = 1.0, p = 0.32). The interaction of timepoint × group did not quite reach significance (F_1,44_ = 3.61, p = 0.064). Taken together, these results suggest that *wv*_*y*_ increased more in the real session than the null session (timepoint × session interaction), and this effect occurred for both force field directions (lack of timepoint × session × group interaction).

**Figure 4.**
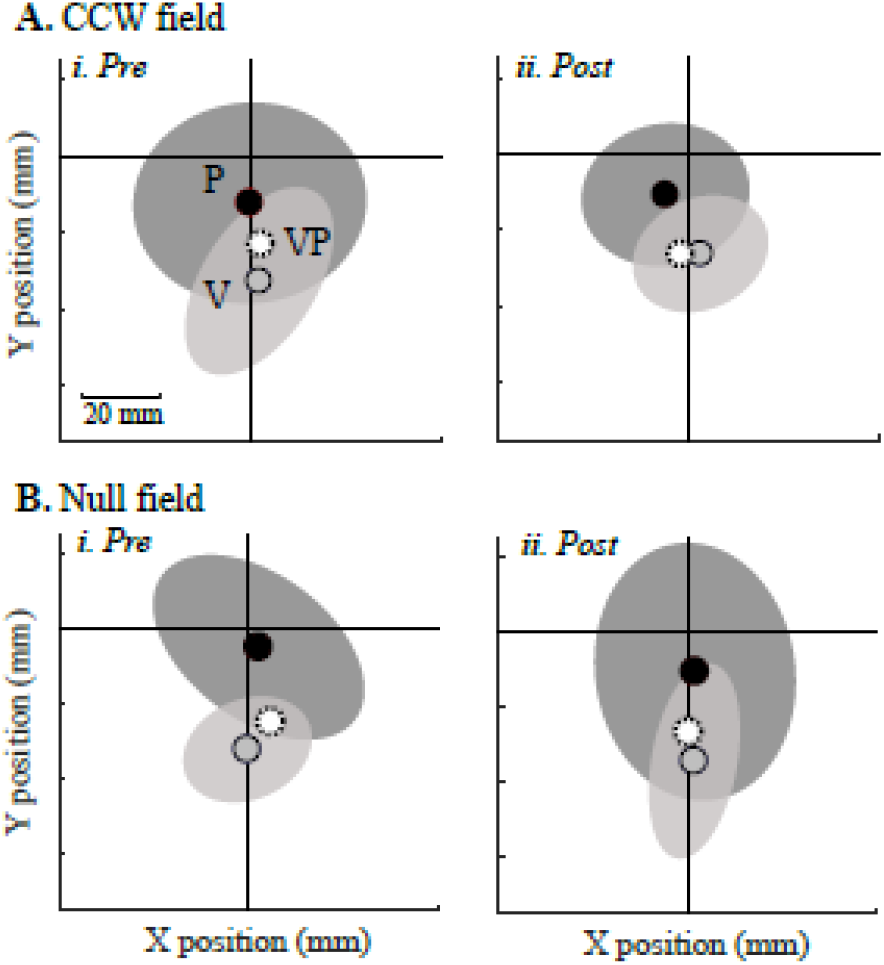
Visuo-proprioceptive estimation data from a subject in the CCW group. Subject was seated in the direction of the negative y-axis, with the target always at the origin. Mean estimates of VP, V, and P targets, with 95% confidence ellipses for the latter two target types. Weight of vision relative to proprioception (*wv*) is computed by comparing relative spatial biases in the three estimates. **A. Real session. i.** Before adapting to the CCW force field, the VP target is estimated midway between the V and P targets in the y-dimension, indicating relatively equal reliance on vision and proprioception (*wv*_*y*_ = 0.53). **ii.** After adapting to the force field, the VP target is estimated closer to the V than the P target in the y-dimension, indicating greater reliance on vision (*wv*_*y*_ = 0.99). In contrast, x-dimension weighting decreased slightly in this session (*wv*_*x*_= 0.57 pre-adaptation and 0.50 post-adaptation). **B. Null session.** On another day, *wv*_*y*_ decreased slightly after an equal number of reaching trials in a null field (0.73 pre and 0.68 post), and *wv*_*x*_ increased (0.48 pre and 0.62 post).

**Figure 5.**
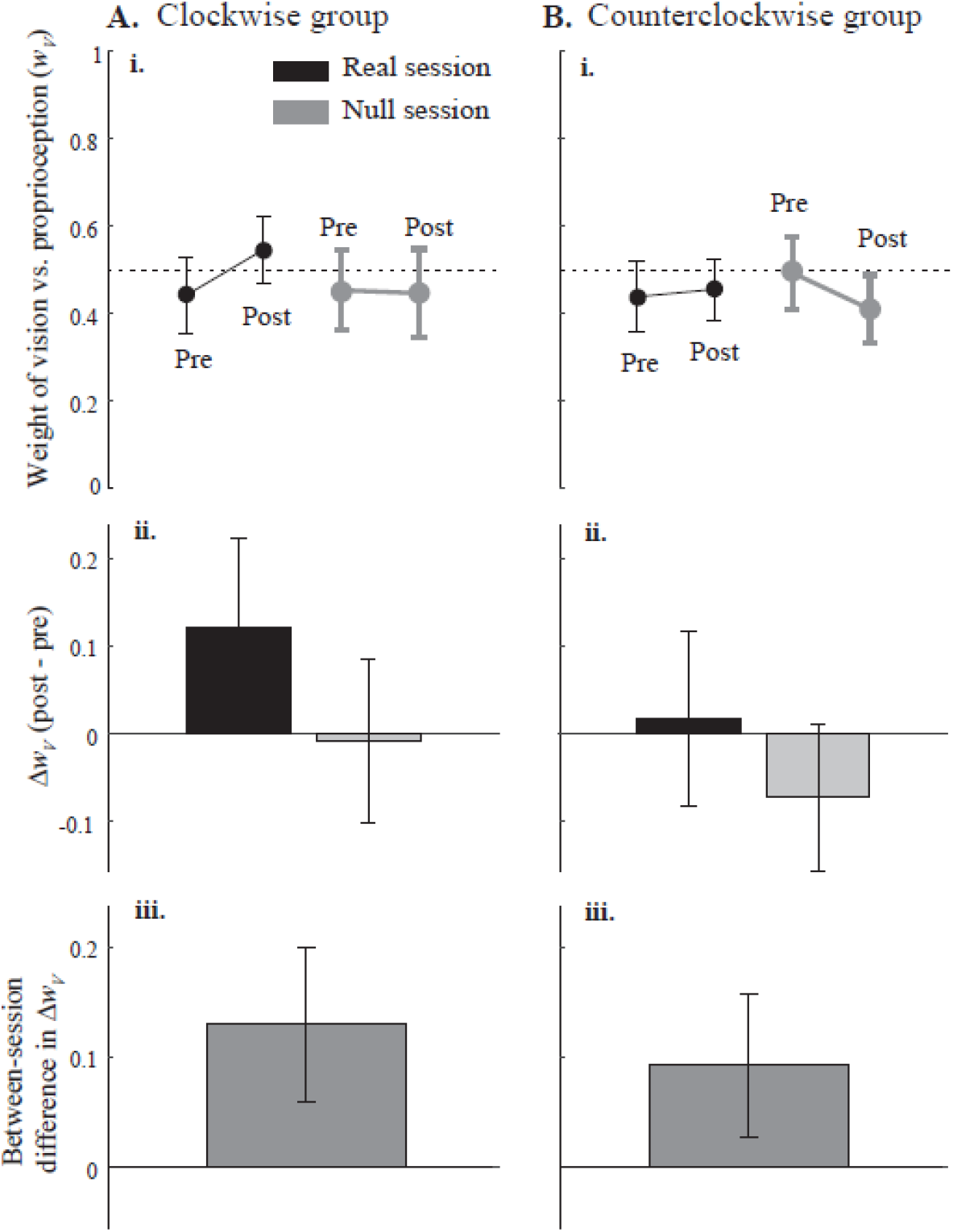
Weight of vision versus proprioception in the y-dimension (*wv*_*y*_) for the clockwise (**A**) and counterclockwise (**B**) groups. **i.** Mean *wv*_*y*_ pre-and post-adaptation block in the real (black) and null (grey) session. 0 corresponds to total reliance on proprioception, and 1 corresponds to total reliance on vision. **ii.** Mean within-session change in *wv*_*y*_. Positive values indicate increased reliance on vision, while negative values indicate increased reliance on proprioception. **iii.** Mean between-session difference in Δ*wv*_*y*_. A significant interaction between session (real, null) and timepoint (pre, post) suggests that *wv*_*y*_ increased after real, but not null, force adaptation, whether the force field was clockwise or counterclockwise. All error bars represent 95% confidence intervals.

Weighting changes in the x-dimension (Fig. 6) were less consistent than in the y-dimension. In the CW group, Δ*wv*_*x*_ was −0.031 ± 0.093 for the real session (mean ± 95% CI) and −0.088 ± 0.079 for the null session. In the CCW group, Δ*wv*_*x*_ was −0.014 ± 0.089 for the real session and −0.049 ± 0.090 for the null session. In other words, on average, subjects reduced their reliance on vision and increased their reliance on proprioception in the x-dimension, regardless of session or group (main effect of timepoint, F_1,44_ = 4.34, p = 0.043). There was no effect of session (F_1,44_ = 1.0, p = 0.32) and no significant interactions (all p > 0.2). However, the groups apparently differed in some way on *wv*_*x*_ (main effect of group, F_1,44_ = 5.82, p = 0.020). Because we have previously observed that subjects vary substantially in weighting of vision vs. proprioception even with no perturbation, we wondered if pre-adaptation *wv*_*x*_ in either of our two groups differed substantially from subjects in earlier studies. To find out, we compared pre-adaptation *wv*_*x*_ in the CW and CCW group with *wv*_*x*_in 80 healthy young adults (mean age 23.4 years) described previously (Liu et al., 2018). Mean *wv*_*x*_ in these three samples was 0.43 ± 0.07, ± 0.05, and 0.49 ± 0.04 (mean ± 95% CI), respectively. A one-way ANOVA found no significant differences among these three samples (F_2,125_ = 1.71, p = 0.18). In other words, although pre-adaptation *wv*_*x*_ favored proprioception more in the CW group than in the CCW group, perhaps explaining the main effect of group on *wv*_*x*_ in the present study, neither sample differed significantly from subjects we have tested previously.

**Figure 6.**
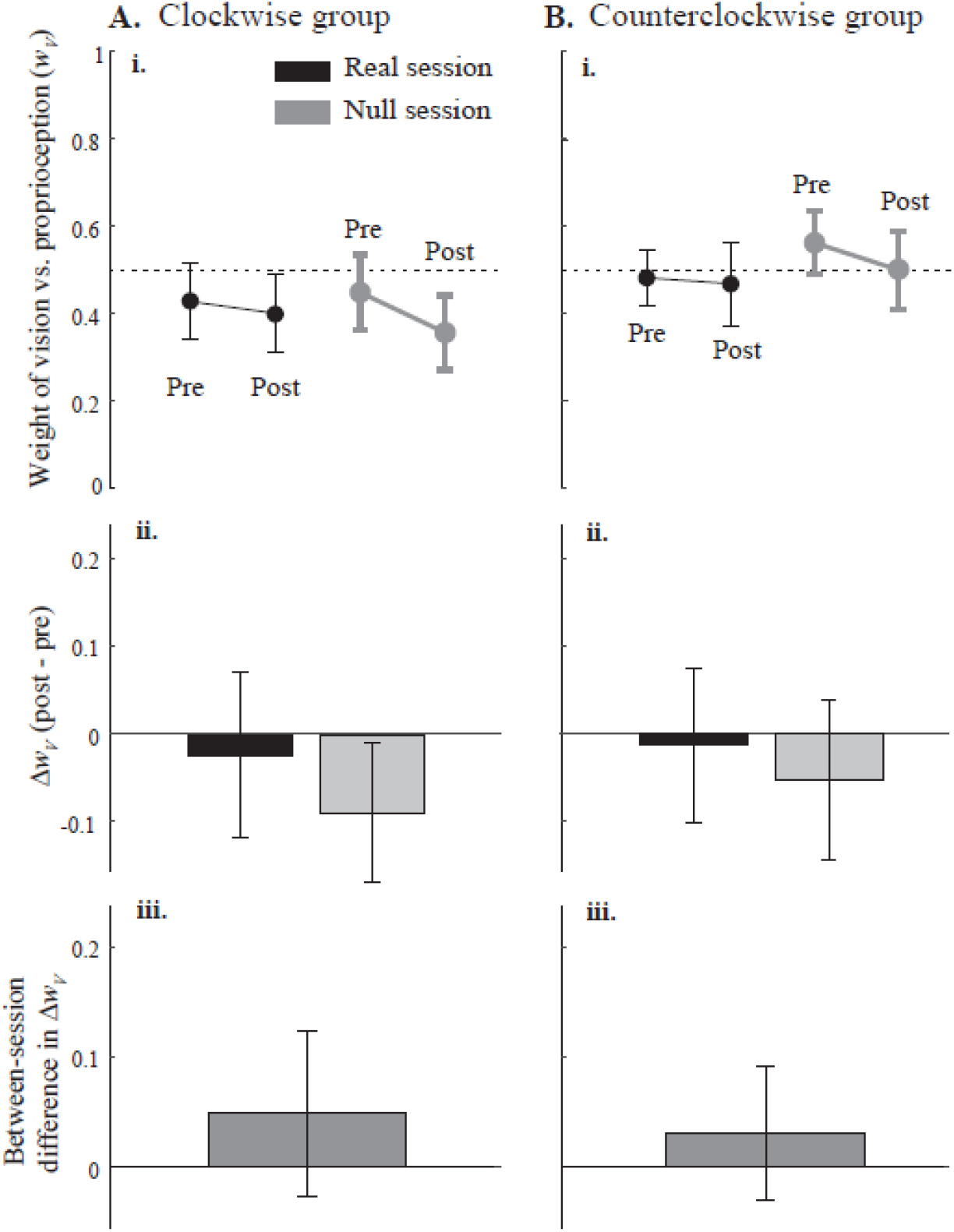
Weight of vision versus proprioception in the x-dimension (*wv*_*x*_) for the clockwiseand counterclockwise (**B**) groups. **i.** Mean *wv*_*x*_ pre-and post-adaptation block in the real (black) and null (grey) session. 0 corresponds to total reliance on proprioception, and 1 corresponds to total reliance on vision. **ii.** Mean within-session change in *wv*_*x*_. Positive values indicate increased reliance on vision, while negative values indicate increased reliance on proprioception. **iii.** Mean between-session difference in Δ*wv*_*x*_. The absence of interaction effects suggests that force adaptation did not significantly affect *wv* in the x-dimension. All error bars represent 95% confidence intervals.

## Discussion

Here we asked whether force field adaptation is associated with changes in visuo-proprioceptive weighting. Subjects increased their weight of vision vs. proprioception in the force field session relative to the null field session, regardless of force field direction, in the straight-ahead dimension. This increase in reliance on vision over proprioception could indicate that the brain interprets the force field as a somatosensory perturbation or sign of proprioceptive unreliability.

### Multisensory processing in motor control

Inherent in any target-directed hand movement is an estimate of hand position, which can be encoded by both vision and proprioception. The brain is thought to weight and combine available sensory estimates to form an integrated multisensory estimate of hand position with which to guide movement (Ghahramani et al., 1997). Multisensory research has made substantial progress in determining the principles by which multisensory integration occurs and demonstrating their relevance in human perception (Angelaki et al., 2009; Fetsch et al., 2013). In addition, it is known that multisensory weights have a role in motor control. E.g., different visuo-proprioceptive weights are evident for different reach target modalities and reach directions (Sober and Sabes, 2005). However, it is not clear whether a perceptual computation such as visuo-proprioceptive weighting is affected by, or plays a role in, motor adaptation or learning.

Motor adaptation is a trial-by-trial process of reducing movement error. E.g., visuomotor adaptation occurs in response to a visual perturbation. A visuomotor rotation paradigm deviates the cursor representing hand position by some angular offset such as 30°. With practice, subjects adapt their movements so the cursor reaches the target. Shifts in proprioceptive estimates of hand position (proprioceptive realignment or recalibration) have been observed in conjunction with visuomotor adaptation (Clayton et al., 2014; Cressman and Henriques, 2009). Force field adaptation involves a somatosensory perturbation rather than visual. When forces are systematically applied to the hand during a reach, e.g. pushing the hand to the right when the subject is trying to reach straight ahead, rightward errors occur at first. However, error feedback recalibrates the sensorimotor map, gradually reducing movement error. Until recently, a force perturbation had been thought to elicit motor adaptation only, as visuo-proprioceptive signals remain veridical (Cressman and Henriques, 2009, 2015); sensory realignment was thought to require inter-sensory misalignment (Sarlegna and Bernier, 2010). However, Ostry et al. (2010) tested arm proprioception and found systematic changes during force adaptation, independent of motor adaptation rate (Mattar et al., 2013). This could reflect a common sensorimotor map, with modifications affecting parameters of multisensory integration.

### Visuo-proprioceptive weighting increased during force field adaptation

To our knowledge, subjects’ weighting of vision vs. proprioception, with both signals available simultaneously, has not been examined in either visuomotor or force field adaptation. We chose to study visuo-proprioceptive weighting in the context of force field adaptation because from the perspective of multisensory processing, the predictions are straightforward: The target is visual, the hand is represented by a visual cursor, and the perturbation disrupting the movement to the target is somatosensory. All these factors would be expected to favor vision over proprioception in the weighting computation. To control for aspects of the task other than the somatosensory perturbation, subjects in the present study performed one session entirely in the null field, and one session with the force perturbation. We predicted that compared to the control session, subjects would up-weight vision relative to proprioception. Our results provide evidence that this is indeed the case. For y-dimension weighting, the presence of a timepoint × session interaction in the absence of a timepoint × session × group interaction indicates that subjects’ increase in weighting of vision was specific to the force field session, regardless of whether that field was CW or CCW. Thus, unlike force field-related spatial recalibration of proprioception (Mattar et al., 2013; D.J. Ostry et al., 2010) or vision (Haith et al., 2008), force field *direction* does not appear to impact visuo-proprioceptive weighting. This may indicate that the force field in general is interpreted by the brain as somatosensory unreliability or perturbation. Literature from postural control supports this conclusion: Decreasing the reliability of somatosensory input increases reliance on the visual and vestibular systems (Mahboobin et al., 2005).

It is important to note that we cannot infer causal relationships from these results. It is possible that force field adaptation, or even exposure to the force field, caused subjects to increase their reliance on vision. However, it is also possible that the observed weighting changes form an unrecognized component of motor adaptation whose absence would impair performance. To distinguish these possibilities, future studies could assess motor adaptation rate, magnitude, and retention after manipulating subjects’ visuo-proprioceptive weighting, perhaps via target modality (Sober and Sabes, 2003) or salience (Block and Bastian, 2010).

### Results were dimension-specific

Interestingly, we found robust evidence of force field-related increase in visuo-proprioceptive weighting for the y-dimension (sagittal plane), but not the x-dimension (lateral plane). We analyzed the x-and y-dimensions separately because of evidence that visuo-proprioceptive weighting computations vary by spatial dimension (van Beers et al., 2002). Certain spatial aspects of motor adaptation are also thought to be controlled separately: Distinct coordinate systems for adaptation of movement direction and extent have been observed (Poh et al., 2017). The force field perturbation in the present study was a curl field, with equal components in the x-and y-dimensions (eq. 2). However, the desired movement was entirely in the y-dimension, since the target was straight ahead of the starting position. In addition, subjects were explicitly instructed to move the manipulandum handle straight ahead. Since visuo-proprioceptive weighting is affected by locus of attention and target salience (Block and Bastian, 2010), these factors in the task design could have created a situation where the straight-ahead dimension was more sensitive to changes in visuo-proprioceptive weighting associated with the force field. This could be tested by altering the dimensional parameters in a new experiment.

An alternative explanation for the lack of force field session-specific change in weighting for the x-dimension could be the weighting characteristics of the two groups. The main effect of group in this parameter may indicate that subjects in the two groups differed in their x-dimension weighting in general, not in a way that changed differently over time or in different sessions. We have observed substantial inter-subject variability in visuo-proprioceptive weighting even in unperturbed situations (Block and Bastian, 2010; Liu et al., 2018). However, the absence of a main effect of session for x-dimension weighting, or any interaction involving session, leads us to hypothesize that even two groups with more similar baseline x-dimension weighting would not change differently in the force field relative to null field session.

One result we did not expect was the near-significant interaction of timepoint × group in the y-dimension. It is indeed possible that this variable changed differently in the two groups. However, any such change would have applied similarly in the force and null sessions. In other words, this interaction, even if truly present, was unrelated to the presence of the force field.

### Considerations for a bimanual sensory estimation task

While similar sensorimotor studies have used this method (Haith et al., 2008; D.J. Ostry et al., 2010), psychometric procedures using two-alternative forced choice (2AFC) tasks are perhaps more common. However, such tasks in motor control experiments have almost always been used to measure proprioceptive spatial alignment, not visuo-proprioceptive integration parameters. A bimanual estimation task presents several advantages over single-modality blocks of 2AFC trials when the goal is to estimate visuo-proprioceptive weighting. Using the left hand, which is not exposed to the force perturbation, to indicate perceived right hand position allows simultaneous estimation in both lateral and sagittal dimensions (Haggard et al., 2000). It is also more analogous to using sensory information for motor planning, and lends itself to mixing up the visual, proprioceptive, and visuo-proprioceptive trial types to make it apparent that the visual and proprioceptive signals, while sometimes presented alone and sometimes together, relate to the same object. Given that multisensory weighting can change instantaneously and differs for different computations, the bimanual approach is better able to assess the computation of interest, multisensory weighting.

A bimanual task in this situation does carry risks. Namely, any intermanual transfer of force field adaptation to the left hand would bias the sensory estimates. However, this risk is small considering that proprioceptive recalibration (Henriques and Cressman, 2012) and motor adaptation do not transfer well to movements with different kinematics or contexts (T. A. Martin et al., 1996), and the movements of the left hand in the sensory task differ in posture, orientation, and movement path from the reaching task. We can reasonably expect that if any transfer were to occur, it should be very small. Indeed, neither group in the present study showed significant aftereffects in the left hand in the force field relative to the null session, suggesting that even within the reaching task, intermanual transfer was absent or negligible.

### Potential neural substrates

Recent years have brought a number of advances in our understanding of the neural substrates of motor adaptation. For example, the early stages of motor adaptation are thought to engage spatial working memory and explicit processes (Seidler and Carson, 2017), associated with activations in DLPFC and inferior parietal lobule (Anguera et al., 2011, 2010; Christou et al., 2016; Taylor et al., 2014). Late adaptation involves implicit processes to a greater extent (Seidler and Carson, 2017). Learning at this point may depend more on the cerebellum (Bernard and Seidler, 2013; Cassady et al., 2018; Flament et al., 1996; Kim et al., 2015). Many neuroimaging and patient studies have also suggested that the cerebellum is critical for motor adaptation (Diedrichsen et al., 2005; T.A. Martin et al., 1996; Tseng et al., 2007; Weiner et al., 1983). Indeed, non-invasive brain stimulation studies have found that manipulating cerebellar excitability can alter the rate of error reduction in motor adaptation (Block and Celnik, 2013; Galea et al., 2011; Jayaram et al., 2012). In addition, plasticity in somatosensory cortex is associated with force adaptation (Ostry and Gribble, 2016); somatosensory training has been found to improve motor adaptation (Darainy et al., 2013; Wong et al., 2012).

Several of the brain regions thought to have a role in motor adaptation are also known to have multisensory visuo-proprioceptive properties, suggesting potential neural substrates for our observation of visuo-proprioceptive weighting changes associated with force field adaptation. First, certain parietal regions have been found to respond to both the “seen” and “felt” position of the limb in monkeys (Graziano, 1999; Graziano et al., 2000), suggesting possible involvement in visuo-proprioceptive integration. Second, while the cerebellum has traditionally been classified as a motor structure and its importance in motor adaptation is well known, this structure also has multimodal sensory responses. Indeed, individual cerebellar granule cells have been found to integrate somatosensory, visual, and auditory inputs (Ishikawa et al., 2015). In humans, the cerebellum has been implicated in multisensory integration for postural control (Helmchen et al., 2017) and reaction time (Ronconi et al., 2017). Finally, the role in perception of regions historically considered unisensory, such as somatosensory cortex, vs. areas considered multisensory, such as portions of PPC, is far from settled (Yau et al., 2015). Such unisensory areas are now known to have multisensory response properties (Ghazanfar and Schroeder, 2006), and likely both modulate, and are modulated by, each other as well as multisensory regions in PPC (Yau et al., 2015). In other words, if changes in visuo-proprioceptive integration accompany force field adaptation, as our results suggest, it is plausible that networks containing regions thought to be involved in both processes (PPC, cerebellum, somatosensory cortex) may mediate the interaction.

## Conclusion

Results of the present study suggest that subjects increase their reliance on vision vs. proprioception when they undergo force field adaptation. This change in visuo-proprioceptive weighting was specific to the sagittal plane, perhaps reflecting the importance of straight-ahead movements in the task design. Force field direction did not play a role, as the effect was comparable for clockwise and counter-clockwise force field exposure. Taken together, these results could indicate that the brain interprets a force field as a somatosensory perturbation and adjusts visuo-proprioceptive integration accordingly.

Author contributions
BMS: Conceptualization, Methodology, Data collection, Data analysis, Writing – original draft.
YL: Methodology, Data analysis, Writing – review & editing.
HJB: Conceptualization, Data analysis, Writing – review & editing, Supervision.

## References

Angelaki, D.E., Gu, Y., DeAngelis, G.C., 2009. Multisensory integration. Curr. Opin. Neurobiol. 19, 452–458. https://doi.org/10.1016/j.conb.2009.06.008

Anguera, J.A., Reuter-Lorenz, P.A., Willingham, D.T., Seidler, R.D., 2011. Failure to engage spatial working memory contributes to age-related declines in visuomotor learning. J. Cogn. Neurosci. 23, 11–25. https://doi.org/10.1162/jocn.2010.21451

Anguera, J.A., Reuter-Lorenz, P.A., Willingham, D.T., Seidler, R.D., 2010. Contributions of spatial working memory to visuomotor learning. J. Cogn. Neurosci. 22, 1917–1930. https://doi.org/10.1162/jocn.2009.21351

Baizer, J.S., Kralj-Hans, I., Glickstein, M., 1999. Cerebellar lesions and prism adaptation in macaque monkeys. J. Neurophysiol. 81, 1960–1965.

Bays, P.M., Wolpert, D.M., 2007. Computational principles of sensorimotor control that minimize uncertainty and variability. J Physiol 578, 387–396.

Bernard, J.A., Seidler, R.D., 2013. Cerebellar contributions to visuomotor adaptation and motor sequence learning: an ALE meta-analysis. Front. Hum. Neurosci. 7. https://doi.org/10.3389/fnhum.2013.00027

Block, H., Bastian, A., Celnik, P., 2013. Virtual lesion of angular gyrus disrupts the relationship between visuoproprioceptive weighting and realignment. J. Cogn. Neurosci. 25, 636–648. https://doi.org/10.1162/jocn_a_00340

Block, H., Celnik, P., 2013. Stimulating the cerebellum affects visuomotor adaptation but not intermanual transfer of learning. Cerebellum Lond. Engl. 12, 781–793. https://doi.org/10.1007/s12311-013-0486-7

Block, H.J., Bastian, A.J., 2012. Cerebellar involvement in motor but not sensory adaptation. Neuropsychologia 50, 1766–1775. https://doi.org/10.1016/j.neuropsychologia.2012.03.034

Block, H.J., Bastian, A.J., 2011. Sensory weighting and realignment: independent compensatory processes. J.Neurophysiol. 106, 59–70.

Block, H.J., Bastian, A.J., 2010. Sensory reweighting in targeted reaching: effects of conscious effort, error history, and target salience. J.Neurophysiol. 103, 206–217.

Cassady, K., Ruitenberg, M., Koppelmans, V., Reuter-Lorenz, P., De Dios, Y., Gadd, N., Wood, S., Riascos Castenada, R., Kofman, I., Bloomberg, J., Mulavara, A., Seidler, R., 2018. Neural predictors of sensorimotor adaptation rate and savings. Hum. Brain Mapp. 39, 1516–1531. https://doi.org/10.1002/hbm.23924

Christou, A.I., Miall, R.C., McNab, F., Galea, J.M., 2016. Individual differences in explicit and implicit visuomotor learning and working memory capacity. Sci. Rep. 6, 36633. https://doi.org/10.1038/srep36633

Clayton, H.A., Cressman, E.K., Henriques, D.Y.P., 2014. The effect of visuomotor adaptation on proprioceptive localization: the contributions of perceptual and motor changes. Exp. Brain Res. 1–14. https://doi.org/10.1007/s00221-014-3896-y

Cressman, E.K., Henriques, D.Y., 2009. Sensory recalibration of hand position following visuomotor adaptation. J. Neurophysiol. 102, 3505–3518.

Cressman, E.K., Henriques, D.Y.P., 2015. Generalization patterns for reach adaptation and proprioceptive recalibration differ after visuomotor learning. J. Neurophysiol. 114, 354–365. https://doi.org/10.1152/jn.00415.2014

Crowe, A., Keessen, W., Kuus, W., van Vliet, R., Zegeling, A., 1987. Proprioceptive accuracy in two dimensions. Percept. Mot. Skills 64, 831–846. https://doi.org/10.2466/pms.1987.64.3.831

Darainy, M., Vahdat, S., Ostry, D.J., 2013. Perceptual learning in sensorimotor adaptation. J. Neurophysiol. 110, 2152–2162. https://doi.org/10.1152/jn.00439.2013

Diedrichsen, J., Verstynen, T., Lehman, S.L., Ivry, R.B., 2005. Cerebellar involvement in anticipating the consequences of self-produced actions during bimanual movements. J.Neurophysiol. 93, 801–812.

Ernst, M.O., Banks, M.S., 2002. Humans integrate visual and haptic information in a statistically optimal fashion. Nature 415, 429–433. https://doi.org/10.1038/415429a

Fetsch, C.R., DeAngelis, G.C., Angelaki, D.E., 2013. Bridging the gap between theories of sensory cue integration and the physiology of multisensory neurons. Nat. Rev. Neurosci. 14. https://doi.org/10.1038/nrn3503

Flament, D., Ellermann, J.M., Kim, S.G., Ugurbil, K., Ebner, T.J., 1996. Functional magnetic resonance imaging of cerebellar activation during the learning of a visuomotor dissociation task. Hum. Brain Mapp. 4, 210–226. https://doi.org/10.1002/hbm.460040302

Foley, J.M., Held, R., 1972. Visually directed pointing as a function of target distance, direction, and available cues. Percept. Psychophys. 12, 263–268. https://doi.org/10.3758/BF03207201

Galea, J.M., Vazquez, A., Pasricha, N., Orban, de, Celnik, P., 2011. Dissociating the Roles of the Cerebellum and Motor Cortex during Adaptive Learning: The Motor Cortex Retains What the Cerebellum Learns. Cereb.Cortex 21, 1761–1770.

Ghahramani, Z., Wolpert, D.M., Jordan, M.I., 1997. Computational models for sensorimotor integration, in: Morasso, P.G., Sanguineti, V. (Eds.), Self-Organization, Computational Maps and Motor Control. North-Holland, Amsterdam, pp. 117–147.

Ghazanfar, A.A., Schroeder, C.E., 2006. Is neocortex essentially multisensory? Trends Cogn Sci 10, 278–285.

Graziano, M.S., 1999. Where is my arm? The relative role of vision and proprioception in the neuronal representation of limb position. Proc.Natl.Acad.Sci.U.S.A 96, 10418–10421.

Graziano, M.S., Cooke, D.F., Taylor, C.S., 2000. Coding the location of the arm by sight. Science 290, 1782–1786.

Haggard, P., Newman, C., Blundell, J., Andrew, H., 2000. The perceived position of the hand in space. Percept. Psychophys. 62, 363–377.

Haith, A., Jackson, C., Miall, R.C., Vijayakumar, S., 2008. Unifying the Sensory and Motor Components of Sensorimotor Adaptation, in: Proceedings of the Neural Information Processing Systems. pp. 1–8.

Helmchen, C., Kirchhoff, J.-B., Göttlich, M., Sprenger, A., 2017. Postural Ataxia in Cerebellar Downbeat Nystagmus: Its Relation to Visual, Proprioceptive and Vestibular Signals and Cerebellar Atrophy. PloS One 12, e0168808. https://doi.org/10.1371/journal.pone.0168808

Henriques, D.Y.P., Cressman, E.K., 2012. Visuomotor adaptation and proprioceptive recalibration. J. Mot. Behav. 44, 435–444. https://doi.org/10.1080/00222895.2012.659232

Ishikawa, T., Shimuta, M., Häusser, M., 2015. Multimodal sensory integration in single cerebellar granule cells in vivo. eLife 4. https://doi.org/10.7554/eLife.12916

Jayaram, G., Tang, B., Pallegadda, R., Vasudevan, E.V.L., Celnik, P., Bastian, A., 2012. Modulating locomotor adaptation with cerebellar stimulation. J. Neurophysiol. 107, 2950–2957. https://doi.org/10.1152/jn.00645.2011

Kim, S., Ogawa, K., Lv, J., Schweighofer, N., Imamizu, H., 2015. Neural Substrates Related to Motor Memory with Multiple Timescales in Sensorimotor Adaptation. PLoS Biol. 13, e1002312. https://doi.org/10.1371/journal.pbio.1002312

Limanowski, J., Blankenburg, F., 2016. Integration of Visual and Proprioceptive Limb Position Information in Human Posterior Parietal, Premotor, and Extrastriate Cortex. J. Neurosci. Off. J. Soc. Neurosci. 36, 2582–2589. https://doi.org/10.1523/JNEUROSCI.3987-15.2016

Liu, Y., Sexton, B.M., Block, H.J., 2018. Spatial bias in estimating the position of visual and proprioceptive targets. J. Neurophysiol. https://doi.org/10.1152/jn.00633.2017

Mahboobin, A., Loughlin, P.J., Redfern, M.S., Sparto, P.J., 2005. Sensory re-weighting in human postural control during moving-scene perturbations. Exp. Brain Res. 167, 260–267. https://doi.org/10.1007/s00221-005-0053-7

Martin, T.A., Keating, J.G., Goodkin, H.P., Bastian, A.J., Thach, W.T., 1996. Throwing while looking through prisms. I. Focal olivocerebellar lesions impair adaptation. Brain 119 (Pt 4), 1183–1198.

Martin, T. A., Keating, J.G., Goodkin, H.P., Bastian, A.J., Thach, W.T., 1996. Throwing while looking through prisms. II. Specificity and storage of multiple gaze-throw calibrations. Brain J. Neurol. 119 (Pt 4), 1199–1211.

Mattar, A.A.G., Darainy, M., Ostry, D.J., 2013. Motor learning and its sensory effects: time course of perceptual change and its presence with gradual introduction of load. J. Neurophysiol. 109, 782–791. https://doi.org/10.1152/jn.00734.2011

Melendez-Calderon, A., Masia, L., Gassert, R., Sandini, G., Burdet, E., 2011. Force field adaptation can be learned using vision in the absence of proprioceptive error. IEEE Trans. Neural Syst. Rehabil. Eng. Publ. IEEE Eng. Med. Biol. Soc. 19, 298–306. https://doi.org/10.1109/TNSRE.2011.2125990

Miall, R.C., Kitchen, N.M., Nam, S.-H., Lefumat, H., Renault, A.G., Ørstavik, K., Cole, J.D., Sarlegna, F.R., 2018. Proprioceptive loss and the perception, control and learning of arm movements in humans: evidence from sensory neuronopathy. Exp. Brain Res. 236, 2137–2155. https://doi.org/10.1007/s00221-018-5289-0

Mon-Williams, M., Wann, J.P., Jenkinson, M., Rushton, K., 1997. Synaesthesia in the normal limb. Proc. Biol.Sci 264, 1007–1010.

Munoz-Rubke, F., Mirdamadi, J.L., Lynch, A.K., Block, H.J., 2017. Modality-specific Changes in Motor Cortex Excitability After Visuo-proprioceptive Realignment. J. Cogn. Neurosci. 1–14. https://doi.org/10.1162/jocn_a_01171

Ostry, D.J., Darainy, M., Mattar, A.A., Wong, J., Gribble, P.L., 2010. Somatosensory plasticity and motor learning. J.Neurosci. 30, 5384–5393.

Ostry, David J., Darainy, M., Mattar, A.A.G., Wong, J., Gribble, P.L., 2010. Somatosensory plasticity and motor learning. J. Neurosci. Off. J. Soc. Neurosci. 30, 5384–5393. https://doi.org/10.1523/JNEUROSCI.4571-09.2010

Ostry, D.J., Gribble, P.L., 2016. Sensory Plasticity in Human Motor Learning. Trends Neurosci. 39, 114–123. https://doi.org/10.1016/j.tins.2015.12.006

Poh, E., Carroll, T.J., de Rugy, A., 2017. Distinct coordinate systems for adaptations of movement direction and extent. J. Neurophysiol. 118, 2670–2686. https://doi.org/10.1152/jn.00326.2016

Ronconi, L., Casartelli, L., Carna, S., Molteni, M., Arrigoni, F., Borgatti, R., 2017. When one is Enough: Impaired Multisensory Integration in Cerebellar Agenesis. Cereb. Cortex N. Y. N 1991 27, 2041–2051. https://doi.org/10.1093/cercor/bhw049

Sarlegna, F.R., Bernier, P.-M., 2010. On the Link between Sensorimotor Adaptation and Sensory Recalibration. J. Neurosci. 30, 11555–11557. https://doi.org/10.1523/JNEUROSCI.3040-10.2010

Sarlegna, F.R., Malfait, N., Bringoux, L., Bourdin, C., Vercher, J.-L., 2010. Force-field adaptation without proprioception: can vision be used to model limb dynamics? Neuropsychologia 48, 60–67. https://doi.org/10.1016/j.neuropsychologia.2009.08.011

Scheidt, R.A., Conditt, M.A., Secco, E.L., Mussa-Ivaldi, F.A., 2005. Interaction of visual and proprioceptive feedback during adaptation of human reaching movements. J. Neurophysiol. 93, 3200–3213.

Seidler, R.D., Carson, R.G., 2017. Sensorimotor Learning: Neurocognitive Mechanisms and Individual Differences. J. Neuroengineering Rehabil. 14, 74. https://doi.org/10.1186/s12984-017-0279-1

Smeets, J.B., van den Dobbelsteen, J.J., de Grave, D.D., van Beers, R.J., Brenner, E., 2006. Sensory integration does not lead to sensory calibration. Proc.Natl.Acad.Sci.U.S.A 103, 18781–18786.

Sober, S.J., Sabes, P.N., 2005. Flexible strategies for sensory integration during motor planning. Nat Neurosci 8, 490–497.

Sober, S.J., Sabes, P.N., 2003. Multisensory integration during motor planning. J Neurosci 23, 6982–6992.

Taylor, J.A., Krakauer, J.W., Ivry, R.B., 2014. Explicit and implicit contributions to learning in a sensorimotor adaptation task. J. Neurosci. Off. J. Soc. Neurosci. 34, 3023–3032. https://doi.org/10.1523/JNEUROSCI.3619-13.2014

Tseng, Y.W., Diedrichsen, J., Krakauer, J.W., Shadmehr, R., Bastian, A.J., 2007. Sensory prediction errors drive cerebellum-dependent adaptation of reaching. J. Neurophysiol. 98, 54–62.

Vahdat, S., Darainy, M., Ostry, D.J., 2014. Structure of Plasticity in Human Sensory and Motor Networks Due to Perceptual Learning. J. Neurosci. 34, 2451–2463. https://doi.org/10.1523/JNEUROSCI.4291-13.2014

van Beers, R.J., Sittig, A.C., Denier van der Gon JJ, 1996. How humans combine simultaneous proprioceptive and visual position information. ExpBrain Res 111, 253–261.

van Beers, R.J., Wolpert, D.M., Haggard, P., 2002. When feeling is more important than seeing in sensorimotor adaptation. Curr.Biol. 12, 834–837.

Warren, D.H., Schmitt, T.L., 1978. On the plasticity of visual-proprioceptive bias effects. J ExpPsycholHumPerceptPerform 4, 302–310.

Weiner, M.J., Hallett, M., Funkenstein, H.H., 1983. Adaptation to lateral displacement of vision in patients with lesions of the central nervous system. Neurology 33, 766–772.

Welch, R.B., Warren, D.H., 1980. Immediate perceptual response to intersensory discrepancy. Psychol.Bull. 88, 638–667.

Wong, J.D., Kistemaker, D.A., Chin, A., Gribble, P.L., 2012. Can proprioceptive training improve motor learning? J. Neurophysiol. 108, 3313–3321. https://doi.org/10.1152/jn.00122.2012

Xerri, C., Merzenich, M.M., Jenkins, W., Santucci, S., 1999. Representational Plasticity in Cortical Area 3b Paralleling Tactual-motor Skill Acquisition in Adult Monkeys. Cereb. Cortex 9, 264–276. https://doi.org/10.1093/cercor/9.3.264

Yau, J.M., DeAngelis, G.C., Angelaki, D.E., 2015. Dissecting neural circuits for multisensory integration and crossmodal processing. Philos. Trans. R. Soc. Lond. B. Biol. Sci. 370, 20140203. https://doi.org/10.1098/rstb.2014.0203

